# Scalable batch-correction approach for integrating large-scale single-cell transcriptomes

**DOI:** 10.1101/2021.12.12.472307

**Authors:** Xilin Shen, Hongru Shen, Dan Wu, Mengyao Feng, Jiani Hu, Jilei Liu, Yichen Yang, Meng Yang, Yang Li, Lei Shi, Kexin Chen, Xiangchun Li

## Abstract

Integration of the evolving large-scale single-cell transcriptomes requires scalable batch-correction approaches. Here we propose a simple batch-correction method that is scalable for integrating super large-scale single-cell transcriptomes from diverse sources. The core idea of the method is encoding batch information of each cell as a trainable parameter and added to its expression profile; subsequently, a contrastive learning approach is used to learn feature representation of the additive expression profile. We demonstrate the scalability of the proposed method by integrating 18 million cells obtained from the Human Cell Atlas. Our benchmark comparisons with current state-of-the-art single-cell integration methods demonstrated that our method could achieve comparable data alignment and cluster preservation. Our study would facilitate the integration of super large-scale single-cell transcriptomes. The source code is available at https://github.com/xilinshen/Fugue.

## Background

Single-cell sequencing offers tremendous opportunities for biomedical research to explore the cellular ecosystem and molecular mechanisms [1]. Advances in single-cell technologies have spurred the establishment of several public repositories of single-cell data, including the Human Cell Atlas (HCA), the Single-cell Expression Atlas and the Mouse Cell Atlas [2, 3]. The HCA project is committed to curate millions to trillions of single-cells for constructing a comprehensive reference map of all human cells. As can be foreseen, integration of super large-scale single-cells across heterogeneous tissues from diverse sources will be a leading wave for deep exploration of biology [4, 5]. Therefore, scalable computational methods are crucial for integration of single-cell transcriptomes and subsequently their translation into biological significance.

Batch effects are fundamental issues to be addressed for integration of single-cell transcriptomes. Batch effects are inevitable as single-cell data were generated by various groups with diverse experimental protocols and sequencing platforms [6]. Considerable progress has been made on batch-correction of single-cell expression. For instance, MNN [7], Scanorama [8] and BBKNN [9] are all based on mutual nearest neighbors (MNNs) identification were successfully applied to guide single-cell integration. The Seurat integration [10] utilizes canonical correlation analysis to identify correlations across datasets and computes MNNs to correct data. Harmony [11] integrates datasets by clustering similar cells from different batches while maximizing the diversity of batches within each cluster. scVI [12] applies a deep learning model to learn shared embedding space among datasets for the elimination of batch effects. However, these methods are not designed for the integration of super large-scale single-cells.

To satisfy this need, we present Fugue, a simple yet efficient solution for batch-correction of super large-scale single-cell transcriptomes. The method extended the deep learning method at the heart of our recently published Miscell approach [13]. Miscell learns representations of single-cell expression profiles through contrastive learning and achieves high performance on canonical single-cell analysis tasks including cell clustering and cell-specific markers inferring. In this study, we expand Miscell through encoding batch information as trainable parameters and adding them into expression profiles. In concept, the gene expression profiles of same cell from different batches could be seen as superposition of the same biological information and different batch information. Fugue incorporates addictive batch information as learnable parameters into gene expression matrix. The batch information can be properly represented after training. By taking batch information as trainable variable, Fugue is scalable in atlasing-scale data integration with fixed memory usage.

We demonstrated the scalability and efficiency of Fugue by applying it to analyze 18 million single-cells obtained from HCA and benchmarked its performance on diverse datasets along with current state-of-the-art methods. We showed that Fugue achieved favorably performance as compared with current state-of-the-art methods. The reference map of HCA dissected by Fugue demonstrated that it can learn smooth embedding for time course trajectory and joint embedding space for immune cells from heterogeneous tissues.

## Results

### Overview of Fugue

Fugue integrates single-cells through adding batch information into expression profile and learns batch information by contrastive learning. Specifically, we construct Fugue as a deep learning-based feature encoder to learn dimension reduction representation of expression profile. Given a set of uncorrected single-cells (**Figure 1A**), Fugue embeds their batch information as a learnable matrix (i.e. batch embedding matrix) and adds them to the corresponding expression profile (**Figure 1B**). A DenseNet of 21 layers [14] is used as feature encoder to learn the additive expression profiles. The feature encoder is trained in a self-supervised manner through contrastive learning (**Figure 1C**) [15]. Contrastive loss minimizes the distance between the cell and its noise-added view, and maximizes the distance between different cells. The trained feature encoder is used to extract feature representations of single-cells (**Figure 1D**). We remove the batch embedding matrix from the input. As a result, only biological signals are retained in the embedding space. The representation could be utilized for downstream analysis such as single-cell cluster delineation (**Figure 1E**). Details are described in **Methods** section.

**Figure 1.**
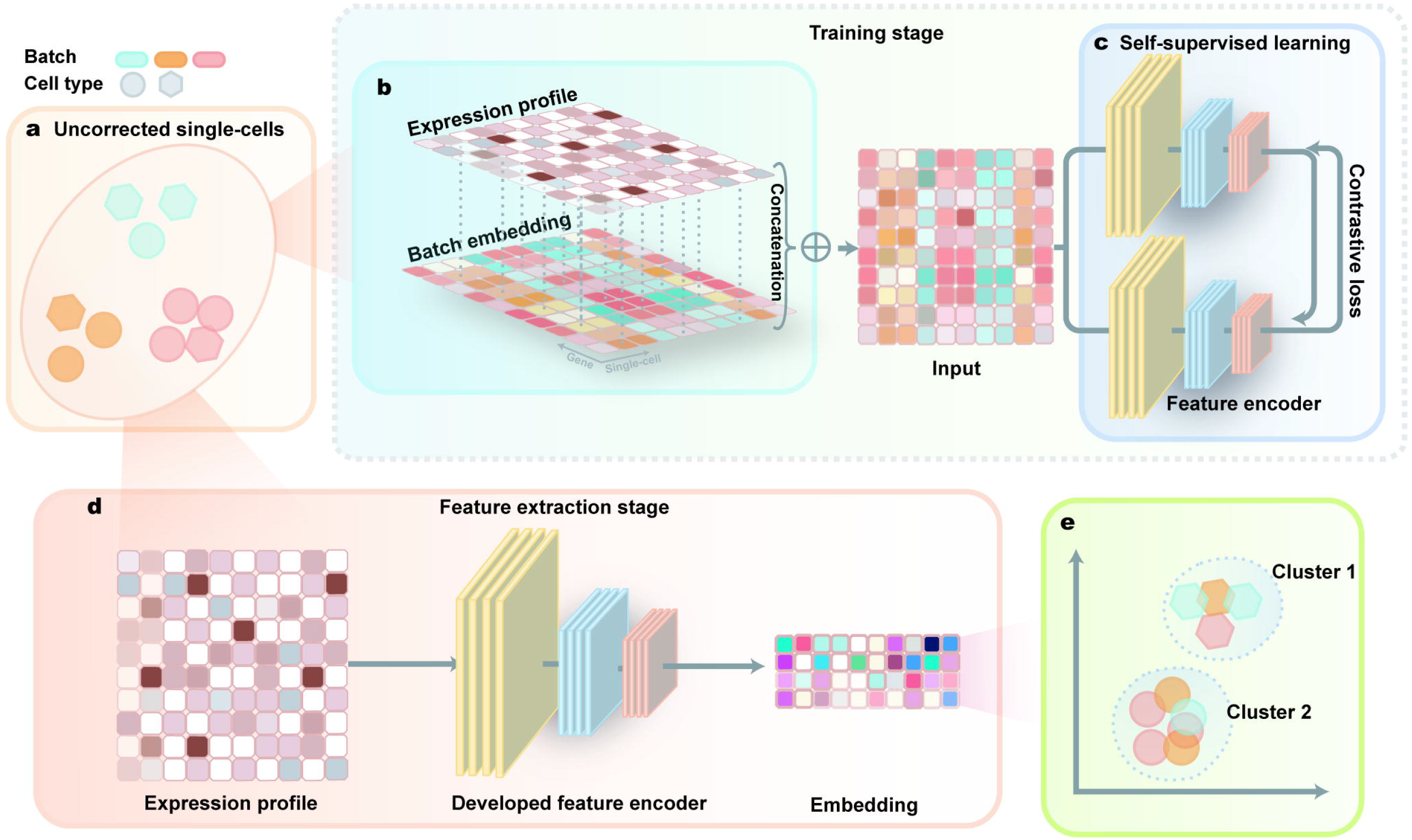
Overview of Fugue. (A) Given a set of uncorrected single-cells, (B) Fugue embedded their batch information as a learnable matrix and added them to the expression profile for feature encoder training. (C) The feature encoder was trained with contrastive loss. (D) At the feature extraction stage, single-cell expression profiles were provided to the feature encoder to extract embedding representation. (E) The embedding representation could be utilized for downstream analysis such as visualization and cell clustering.

### Benchmark evaluations

On the *simulation dataset* of 30,000 cells of 3 cell types among 5 batches, each cell type was divided by batches (**Figure 2A**) before batch correction. After integration with Fugue, cells of the same types were well-mixed and cells of different types were dispersed across batches (**Figure 2B**). In addition, we ran Fugue on this simulation dataset after removing a specific cell type from four batches (See **Methods and Supplementary Figure 1**). The result showed that Fugue could maintain batch-specific cell types (**Figure 2C**).

**Figure 2.**
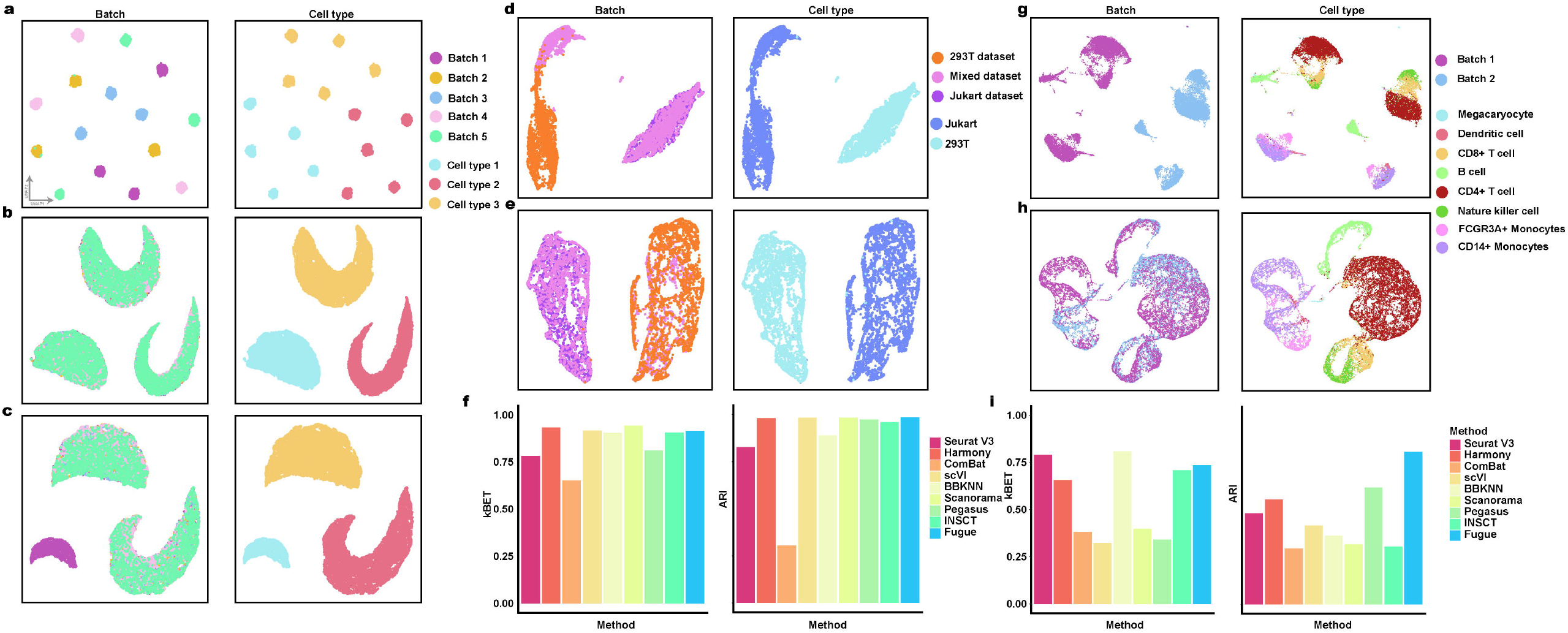
Benchmark of batch-correction performance of Fugue across the *simulation, cell line* and *PBMC datasets*. (A) UMAP plot of cells from simulation dataset, which consists of 5 different batches and 3 cell types. (B) UMAP visualization of Fugue batch effect removing performance on the *simulation dataset*. (C) UMAP plot of Fugue batch effect removing performance on the *simulation_rm* dataset. (D) UMAP plot displays cells from the *cell line dataset*, which consists of 3 different batches and 2 cell types. (E) UMAP plot of Fugue batch effect removing performance on *cell line dataset*. (F) Quantitative assessments of different batch effect removal methods on *cell line dataset*. (G) UMAP plot displaying cells from *PBMC* dataset, which consists of 2 different batches and 8 cell types. (H) UMAP plot of Fugue batch effect removing performance on *PBMC* dataset. (I) Quantitative assessments of different batch effect removing methods on *PBMC* dataset. For (A-E, G-H), cells are colored by batch (left panel) and cell type (right panel).

We used this simulation dataset to search for three hyperparameters that are adjusted for contrastive learning, including size of memory bank and momentum coefficient. The kBET [16] and ARI scores were applied to evaluate its performance (**see Methods**). Fugue was insensitive to variation of these hyperparameters in terms of ARI and kBET scores (**Supplementary Figure 1A and 1B**). Data augmentations include random dropout and position shuffling. We set the dropout rate to 30% and random shuffle rate to 10% based on the value of ARI and kBET (**Supplementary Figure 1C**).

We compared Fugue to 8 single-cell integration methods on the *cell line* (n = 9,531) and *PBMC* (n = 28,541) *datasets* (See Methods, **Figure 2D, G**). Fugue yielded similar result as these 8 methods on UMAP plots (**Figure 2E, H and Supplementary Figure 2**,**3**). Quantitatively, Fugue achieved comparable kBET and ARI scores (**Figure 2F, I**).

### Fugue could accurately remove batch effects

We applied Fugue to integrate all available data from HCA repository (75 cohorts totaling 18,056,192 cells) (**Supplementary Table 1**). The batch effect removing efficiency of Fugue was evaluated on three datasets included in HCA, including the *census of immune project*, the *lung and* the *brain dataset*.

Common cell types of the *census of immune project* (cord blood, n = 133,264; bone marrow, n = 176,571) revealed a minimal overlap before integration (**Supplementary Figure 4A**). Fugue clustered cells into biologically coherent groups and removed batch-specific variations (**Figure 3A**), and UMAP plot was similar to the aforementioned benchmark methods (**Supplementary Figure 4B-H**). Fugue achieved comparable kBET and ARI scores as compared with these methods (**Figure 3B**).

**Figure 3.**
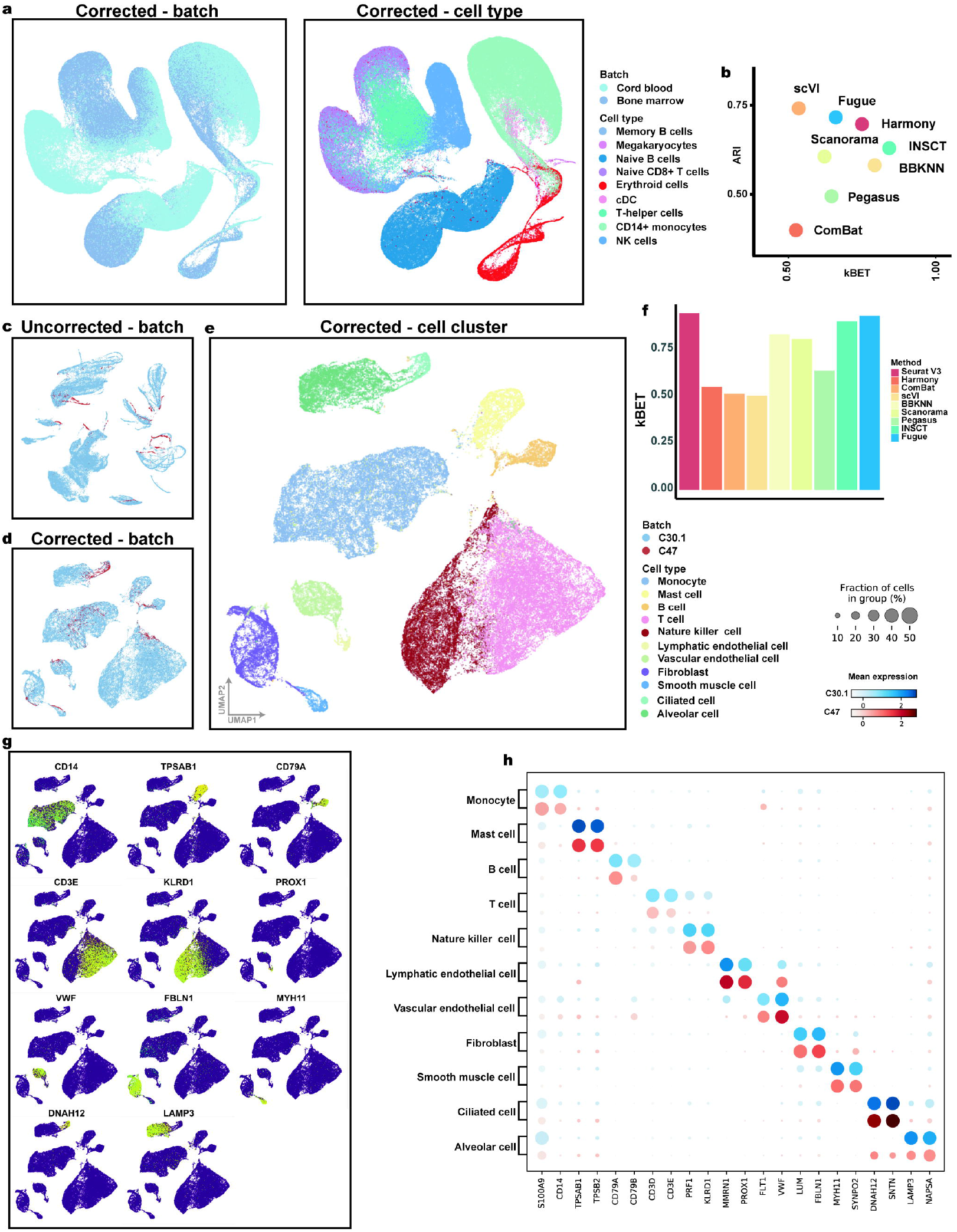
Assessment of the batch-correction performance of Fugue. (A) UMAP plot of Fugue batch effect removing performance on the *census of immune project*. Cells are colored by batch in the left panel and cell ontology label provided in the original publication in the right panel. (B) Quantitative assessments of different batch effect removal methods on the *census of immune project*. (C-E) UMAP plot depicting cells in the *lung dataset* before (C) and after (D-E) Fugue integration. Cells are colored by batch in (C-D) and cell cluster label in (E). (F) Bar plot depicting kBET scores of different batch effect removing methods on the *lung dataset*. (G) Expression of cell type markers across the feature embedding space. Dark and light colors represent low and high relative expression values, respectively. (H) Dot plot representing cell markers across batches. The size of each circle reflects the percentage of cells in a cluster where the gene is detected, and the color intensity reflects the average expression level within each cluster.

The C30.1 (n = 75,387) and C47 (n = 2,532) from *lung dataset* showed minimal overlap before batch correction (**Figure 3C**). After correction with Fugue, cells from different datasets were mapped into corresponding area (**Figure 3D**). The UMAP plot was consistent with the aforementioned benchmark methods (**Supplementary Figure 5**). Quantitatively, Fugue achieved comparable kBET and ARI scores with these methods (**Figure 3F**). Unsupervised clustering and cell types annotation revealed 11 cell types in the *lung dataset*, including monocytes, mast cells and ciliated cells (**Figure 3E**). Conventional cell markers [17, 18] were expressed uniquely in each cell cluster (**Figure 3G**), and invariant across batches (**Figure 3H**).

We evaluated the batch removing efficiency on the *brain dataset* with the same process applied for the *lung dataset*. The result also demonstrated that Fugue can robustly integrate cells from multiple studies (**Supplementary Figure 6**).

### Fugue captures the real batch information

We hypothesize that sequencing samples of the same cohort are subjected to lower batch variation. Therefore, the batch embeddings of samples from the same cohort should be more similar than those from different cohorts.

We extracted the batch embeddings of 373 samples from 75 cohorts in HCA. The result showed that samples from the same cohort had higher similarity of batch embeddings as compared with samples from different cohorts (**Supplementary Figure 7A)**. We found that batch embeddings of 4 patients from the C2 cohort are almost identical (**Supplementary Figure 7B**), which was consistent with the previous report that there was no batch effect among these four patients [19]. For the *census of immune project*, we observed higher similarity of batch embeddings within the same batch than between batches (**Supplementary Figure 7C**). For the PBMC and tonsil tissue from C39 subjected to the same sequencing protocol [20], we also observed high similarity among them, especially among samples from the same tissue (**Supplementary Figure 7D**).

### Fugue aligned precise immune cell subtypes in HCA

Immune cells are highly homogeneous across tissues [21]. Therefore, Fugue should be able to map the same immune cell types together across HCA. Forty-six clusters were inferred from HCA (n = 3,424,607) (**Supplementary Figure 8A and Supplementary Table 2**). Most clusters consist of multiple cohorts, while some come from specific organs (**Supplementary Figure 8B**). For example, 26 projects had over 100 cells in endothelial cell_1 cluster; C2 was the only project associated with lymphatic tissue [19] and made up the majority of lymphatic endothelial cell cluster (endothelial cell_7) (**Supplementary Figure 8B**).

We reclustered the immune cells **(Supplementary Figure 9)** corresponding to 17 subtypes (**Figure 4A**). Different cell types were readily separable from each other, and dataset specific cell types were retained, such as *in-vivo* stimulated NKT cells (**Figure 4A**). Canonical markers were expressed in corresponding cell types (**Figure 4B**). For example, Pan-B cell markers *CD79A* and *CD79B* were expressed in B cell clusters. B cell precursor specific markers *VPREB1* and *IGLL1* were expressed in the relevant cell type. We observed stable expression of marker genes among batches (**Figure 4C, D and Supplementary Figure 10**). For example, natural killer cell markers *PRF1* and *KLRD1* were expressed in all of the 27 cohorts (**Figure 4D**).

**Figure 4.**
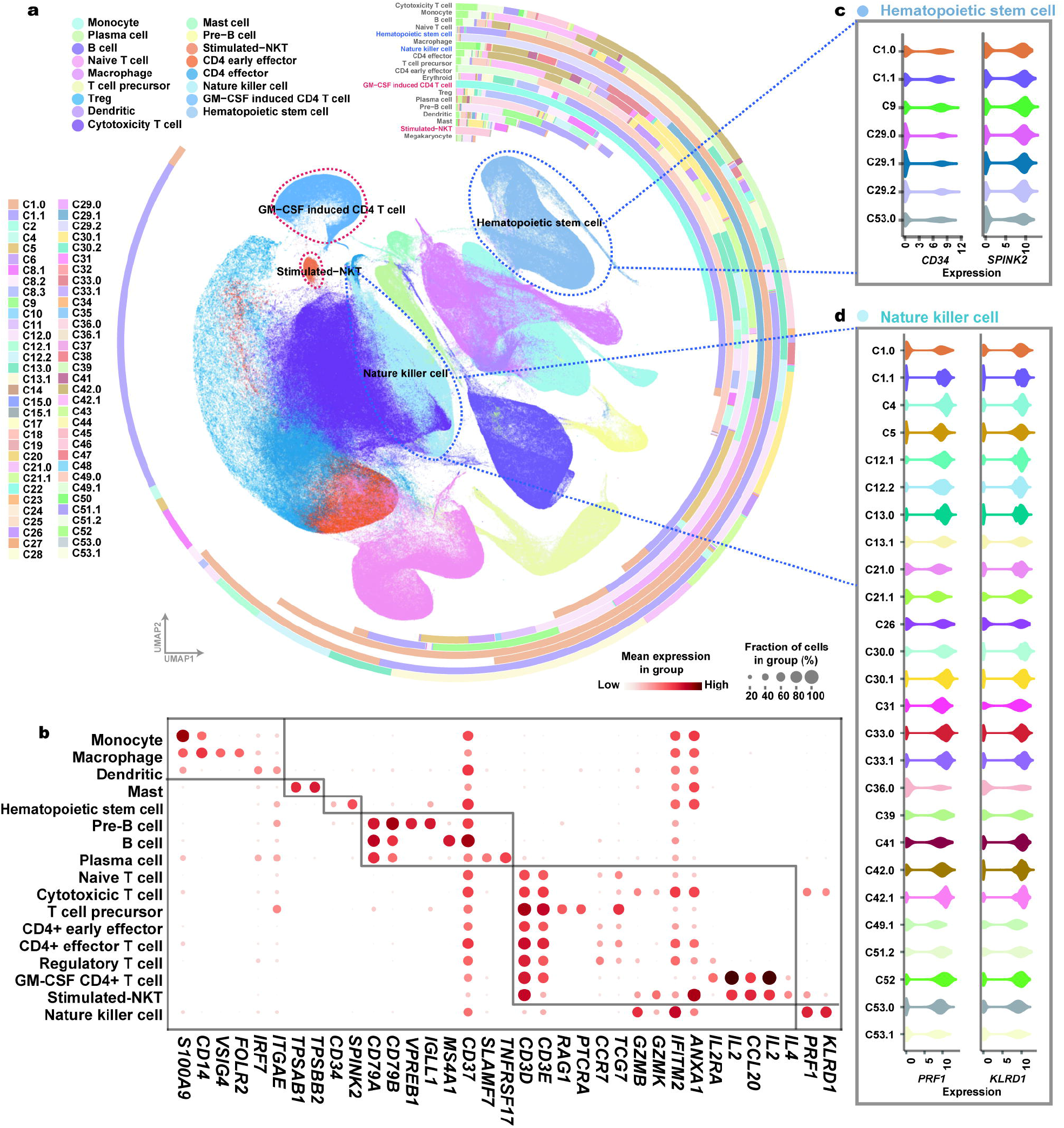
Joint analysis of all immune cells across HCA repository with Fugue. (A) UMAP plot of the 17 immune cell types inferred from Fugue. Cells are colored by cell type labels. (B) Dot plot showing cell type markers across cell clusters. The size of each circle reflects the percentage of cells in a cluster where the gene is detected, and the color intensity reflects the average expression level within each cluster. (C-D) Violin plot deciphering expression levels of cell type markers for hematopoietic stem cells (C) and natural killer cells (D) across HCA cohorts. Cohorts with more than 1000 cells in each cluster were displayed.

### Fugue integrates time course development trajectories

On the *embryonic mouse cardiac dataset*, batch effects were observed among embryo development stage before correction (**Figure 5A**). Fugue integrated cells from different embryo periods (**Figure 5B**). We classified the single-cells into 5 cell types based on the specific cell markers (**Figure 5C, D and Supplementary Figure 11**). The FLE dimension reduction showed that expression representation extracted from Fugue captured the embryonic developmental trajectories for each cell type (**Figure 5E**). Cells from embryonic (E) day 10.5, 13.5 and 16.5 were orderly arranged according to pseudo-time trajectory (**Figure 5E**). The expression patterns of canonical cell differentiation markers were consistent with developmental stages (**Supplementary Figure 12**). For instance, early erythrocyte markers *GYPA* and *TFRC* expressed highly in E10.5 erythrocytes and negatively correlated with pseudo-time (**Supplementary Figure 12**), which was consistent with the previous studies [22, 23].

**Figure 5.**
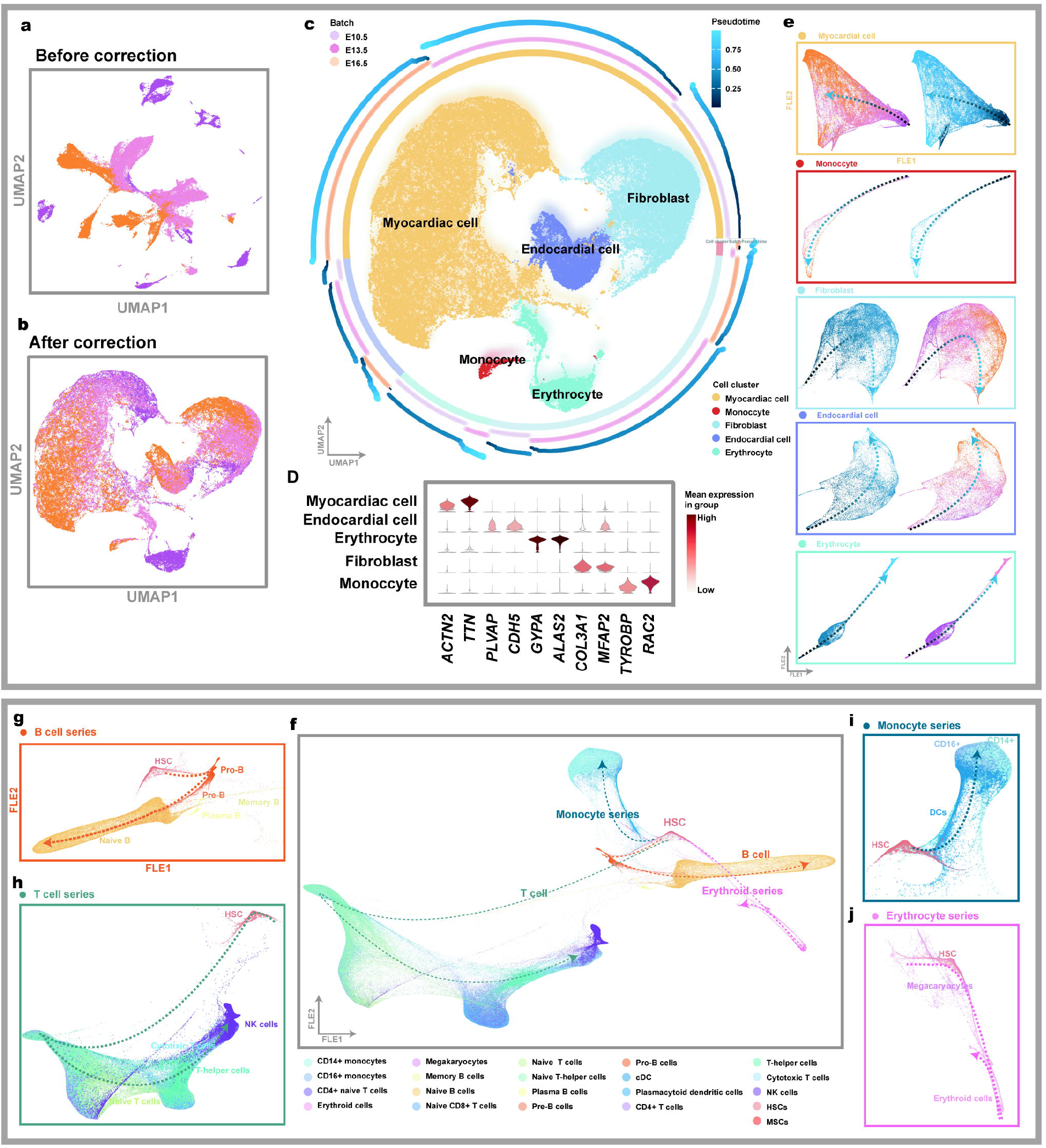
Joint analysis of Fugue on batch-correction and gene expression trajectory recovering during cell development. (A-B) UMAP plot of cells from the *embryonic mouse cardiac dataset* before (A) and after (B) Fugue integration. Cells are colored by batch. (C) UMAP plot of the *embryonic mouse cardiac dataset* integrated by Fugue, colored by cell clusters. The surrounding circle plot from inner to outer shows the cell types, batch labels and pseudo-time scores of the 1% randomly downsampled cells. (D) Violin plot deciphering expression levels of cell type markers across cell clusters. The color intensity reflects the average expression level within each cluster. (E) FLE plot revealing time course trajectories of cardiac development across different cell types. Arrows indicate inferred cell state transition directions from early to late pseudo time. (F) FLE plot revealing cell state transition directions from HSC to all main blood lineages. (G-J) FLE plots of the main development trajectory from HSC to B cell, T cell, monocyte and erythrocyte, respectively. B cell series (G) were separation from HSCs towards B cell progenitors, precursors of B cells and matured naïve B cells. B cells also differentiate into mature B cells, plasma cells and memory B cells. T cell series trajectory (H) was started from HSCs, followed by naïve T cells and finally mature T cells and NK cells. Monocytes series trajectory (I) was started from HCA, and transferred into DCs and CD14+ and CD16+ mature monocytes. Erythrocyte series (J) differentiates from HSCs to megakaryocytes and erythroid cells. Pro, progenitor; Pre, precursor; HSC, hematopoietic stem progenitor cell; DC, dendritic cell; cDC, canonical dendritic cell; NK cells, natural killer cell; MSC, multipotent progenitor cell.

We next recovered cell development trajectories during hematopoiesis from the *census of immune project*. FLE plot indicated clear overlap of the identified cell types from cord blood and bone marrow across pseudo-time trajectory (**Supplementary Figure 13**). Clear trajectories that quadrifurcate from hematopoietic stem cells (HSCs) into B cell, T cell, mono-dendritic and megakaryocyte-erythroid series were constructed (**Figure 5F**). The trajectories were ordered by cell development stages and branching by cell differentiation types (**Figure 5G-J**).

## Discussion and conclusions

In this study, we attempt to tackle the batch effect removal issue in the rapidly developing single-cell transcriptomic with a simple yet effective solution. Fugue could be deployed as a scalable deep learning model to integrate single-cells of any magnitude with fixed memory. We provide evidence that Fugue showcases superior performance in terms of integrating millions of single-cells from various sources. Fugue is expected to assemble all human cells to construct a comprehensive single-cell atlas.

In application, we showcase the robustness of Fugue in super large-scale datasets integration. Specifically, Fugue was applied to integrate all available single-cells among HCA repository. Three datasets included in HCA were utilized to represent the data integrated effectiveness of Fugue, for that most of the benchmark methods cannot handle atlasing-scale datasets due to memory overflow. Moreover, there are currently no suitable indicators to assess the batch-correction performance of complex datasets with multiple distinct or dataset-specific cell types spanning dozens of batches. Fugue performed on par with current state-of-the-art single-cell integration methods in terms of batch-correction and cluster preservation performance. Fugue therefore offers better trade-offs between data integration performance and scalability, and it is a key advantage of Fugue to integrate super large-scale datasets. Furthermore, we show that Fugue could integrate millions of immune cells to reflect delicate cell functional status while retaining distinct cell subtypes. Additional analysis demonstrates that time course trajectories could be correctly constructed and ordered after single-cell integration by Fugue. The algorithm can thus facilitates the exploration of subtle biological differences among atlasing-scale datasets.

A great deal of batch-correction methods learn batch information based on prior assumptions. For example, Combat assumes batches as a function of gene expressions [24]. Methods based on MNN learn batch information through paired cells between batches, and highly depend on the qualities of MNNs [7, 8]. Fugue is a hypothesis-free deep learning network. It simply learns batch information through contrastive learning and does not require domain specific knowledge. The flexibility of this approach could be demonstrated through the integration of single-cells from HCA. For that explicit batch information are not always available from researchers, we employed sample labels as batch information for HCA projects. We demonstrated the compatibility of this configuration through benchmark the performance of Fugue with current state-of-the-art methods, for which accurate batch labels were set. The batch information learned by Fugue also show little variation within the same batch as compared with that between batches (**Supplementary Figure 7**). Therefore, the simple batch correction approach is flexible and can be a good candidate for multi-millions of single-cells where explicit batch information are not always available.

Although immune cell markers have been studied extensively, the knowledge might be limited by their definition via a restricted set of organs or cell types. The integrated analysis of atlasing-scale single-cells enabled cross-organ comparisons and provide new perspectives for the understanding of marker genes. Based on the reference map of HCA, we found many conventional immune cell markers are expressed in nonimmune cell types. For example, conventional monocyte marker *S100A9* was expressed in esophageal squamous epithelium cells (Epithelial cell_4) (**Supplementary Figure 9**), which was confirmed by previous studies [25, 26]. Canonical HSC marker *SPINK2* was expressed higher in epididymal epithelial cells (Epithelial cell_7) than HSCs (Stem cell_1) (**Supplementary Figure 9**). The enrichment of *SPINK2* in epididymal tissue was confirmed in the previous report [27].

Fugue could be improved in several aspects. First, as an artificial intelligence model, black-box nature of the approach is a limitation that should be resolved [28, 29]. We explored the batch embedding matrix and found that similar batches have more similar batch embeddings than dissimilar batches. It brings insights into the interpretability of batch information learned by Fugue. Second, as an unsupervised learning model, hyperparameters tuning might to some extent influence the performance of Fugue [30]. In our analysis, we proved the stability of Fugue to hyperparameters tuning (**Supplementary Figure 1**). We also used the same hyperparameters of model structure throughout the study to ensure the generalization of the result.

In summary, we present Fugue, a simple yet efficient deep learning model for super large-scale single-cell transcriptomes integration. We anticipate Fugue will be helpful for researchers to transform growing scale of single-cell transcriptomes into the understanding of biology and disease, driving new ways for disease diagnosis and treatment.

## Methods

### Batch embedding

The key idea of batch-effect removal is decoupling biological signals from nuisance factors of batch effects. We explicitly encoded batch information as a learnable batch embedding matrix (*BE*) and added them to expression matrix (*E*) to obtain expression matrix with batch embedding information (*X* = *BE* + *E*), subsequently performing feature representation learning on *X*. The batch embedding matrix *BE* was randomly initialized and updated during training. For the purpose of point-wise addition between BE and E, the dimension of matrices BE and E must be identical.

### Network architecture and training

We used DenseNet architecture [14] as feature encoder to learn expression embedding of single-cells. The DenseNet has 21 layers that are consisted of 4 dense blocks. The DenseNet architecture is featured by concatenating all the outputs from preceding layers as input for the next layer to make feature transmission more efficient. We replaced convolutional layer of the DenseNet with linear layer to make it able to process gene expression matrix. Self-supervised learning with momentum contrast [15] was adopted to train the feature encoder. We applied multi-layer perceptron (MLP) as project head, which was demonstrated to be beneficial for contrastive learning [31].

Here we adapt contrastive learning for feature encoder development, through which the model was trained by constructing positive and negative pairs [32]. For a given integrated input *I*_*cell*_, a feature encoder represents it as *C*_*q*_ *= f*_*q*_ *(I*_*cell*_), where *f*_*q*_ *is a* query encoder network and *C*_*q*_ is a query sample. A key encoder network *f*_*k*_ encode the noise-adding view of the input *I*_*cell+*_ *as C*_*k+*_ (likewise, *C*_*k+*_ *= f*_*q*_ *(I*_*cell+*_)). One cell *C*_*q*_ and its noise-adding view *C*_*k+*_ form a positive pair, and assemble with a different cell *C*_*k-*_ *to* form a negative pair. The contrastive loss is optimized through learning the same representation of the positive pairs and dissimilar representation of negative pairs:

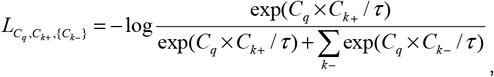

where *C*_*k-*_ denotes a dictionary of the negative samples. The dictionary was built as a queue *C*_*k1-*_, *C*_*k2-*_, …, *C*_*kn-*_. The current mini-batch en queue and the oldest mini-batch de queue. We set the queue size to 10% of the training data. τ is a temperature hyper-parameter and was set to 0.2. We performed data augmentations through random zero out to 30% and shuffling to 10% of genes. These hyperparameters were determined through grid search (see **Supplementary Figure 1**).

The parameters of query encoder □_*q*_ were updated by back-propagation; the parameters of key encoder □_*k*_ were updated according to □_*q*_:

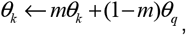

where *m* stands a momentum coefficient, which was set to 0.999. We trained the network at a learning rate of 0.01. The training was ended until loss did not improve over a specified number of epochs (see **Supplementary Table 3**).

The network was trained through mini-batch stochastic gradient descent algorithm [33] with a weight decay of 1e-4. The convergence speed of deep learning model is affected by batch-size [34]. For that we integrated thousands to multi-millions single-cells, we set the size of mini-batch from 16 to 256, which was dependent on the volume of training data (**Supplementary Table 3**).

At the stage of feature extraction, we applied the developed feature encoder *f*_*q*_ as feature extractor. Only expression matrix *E*_*cell*_ was provided to *f*_*q*_:

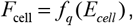

where *F*_*cell*_ is the feature representation of the single-cell transcriptome.

### Data sources

#### Simulation dataset

We simulated a total of 30,000 single-cell read counts using Splatter package[35]. The resultant *simulation dataset* contains 3 cell types; each cell type consists of 5 batches (**Figure 2A**). Each batch contains 2000 genes with a differential expression factor of 0.4. To estimate the performance of Fugue on batch-specific cell types detection, we manually removed cell type 1 from batches 2-5 and maintained them in batch 1 (**Supplementary Figure 14**). We named the resultant dataset as *simulation_rm dataset*.

#### Cell line dataset

This dataset consists of the cell lines of “Jurkat”, “293 T” and the 50/50 mixture of both cell lines [36]. The dataset is composed of 9,531 single-cells generated by 10x 3’ protocol. For mixture cell lines, cells were clustered with Louvain algorithm based on Scanpy pipeline. Cell clusters with high expression of *XIST* were annotated as “293 T” while others as “Jurkat”.

#### Human peripheral blood mononuclear cell (PBMC) dataset

The data included two batches of PBMC from five samples [37]. One sample was excluded from the analysis because it was stimulated *in vitro*. This dataset contains 28,541 single-cells, which could be grouped into B cells, CD4+ T cell, CD8+ T cell, NK cells, monocytes, megakaryocytes and dendritic cells. The cell labels were provided by the original publication [37].

#### Human cell atlas

We downloaded single-cell data from HCA portal [2] on 16 July 2021. Fifty-three projects (C1-C53) following HCA data processing pipeline were collected (**Supplementary Table 1**). These projects consist of 75 cohorts. We filtered out samples with available cell numbers less than 1000. A total number of 373 samples were maintained for downstream analysis. This dataset contains 18,056,192 cells from multiple organs, including blood, lung, brain and cardiac (**Supplementary Table 1**). We used the sample labels as batch information given that explicit batch information is not always available for every dataset. All of that 18,056,192 cells were used by Fugue for batch information learning. A total of 3,424,607 cells with more than 500 expressed genes were utilized to construct the reference map of HCA.

The following projects were selected for the assessment of dataset alignment and biological significance preservation performance of the reference map. The first dataset was the c*ensus of immune project* (C1). The c*ensus of immune project* consists of two batches that can be referred to as cord blood (C1.0) and bone marrow (C1.1). The two batches contain immune cells from diverse development statuses. We downloaded cell type labels from HCA repository on 28 August 2020. The *lung dataset* consists of C30.1 and C47. Both C30.1 and C47 came from lung tissue and have similar cell types [18, 38]. The *brain dataset* consists of C19, C28 and C32, which contain cells from different subsections of brain tissue with overlapping cell types among each other [17, 39, 40]. The *embryonic mouse cardiac dataset* consists of C18 and C20. C18 contains mouse cardiac cells from embryonic state of E10.5 and E13.5. C20 includes mouse cardiac cells from embryonic state of E16.5. Only healthy embryos were taken into account in this analysis. Since the original author of C18 denotes batch effect exists between cells from E10.5 and E13.5 mouse [41], the embryonic periods were employed as batch labels for the benchmarking methods.

### Data prepossessing

We applied Scanpy (**version 1.7.0**) for data preprocessing. We used “highly_variable_genes” function with the default parameters to identify highly variable genes. A total of 1,959 and 2,085 HVGs were selected from the *cell line* and *PBMC* datasets, respectively. For HCA project, 14550 genes shared among datasets were selected.

For all of the aforementioned datasets, we normalized the count matrix to counts per million normalization (CPM) and took logarithmic transformation (i.e. log2(CPM+1)). Subsequently, the expression of each gene was scaled by subtracting its average expression then divided by its standard deviation. The scaled expression matrix was applied as inputs for the model.

### Benchmark methods

We benchmarked the performance of Fugue with eight state-of-the-art batch correction methods, including Seurat V3, ComBat, Harmony, BBKNN, Scanorama, scVI, Pegasus L/S adjustment and INSCT. All methods were performed with the default parameters (see **Supplementary Table 4** for detailed information) throughout the study. Seurat V3 ran out of memory on our server (maximum memory: 256 Gb) for dataset with more than 100,000 cells and therefore it was not evaluated on dataset >100,000 cells. For the c*ensus of immune project*, cord blood and bone marrow were utilized as batch information. We employed different cohorts as batch information for the *lung* and *brain datasets* and embryonic development periods as batch information for the *embryonic mouse cardiac dataset*. We provided these methods with explicit batch information because it’s the general configuration and suitable for these methods [7-12].

### Evaluation functions

We employed kBET acceptance rate [16] for the assessment of batch effect through *Pegasus* package [42]. The kBET acceptance rate measures whether batches are well-mixed in the local neighborhood of each cell. The resulting score ranges from 0 to 1, where a higher score means a better mix. We computed kBET scores based on each cell type and used the average score to evaluate the degree of batch mixing. The adjusted rand index (ARI) score was applied to evaluate batch correction method in terms of cell type mixing. The ARI score measures the percentage of matches between two label lists. The resulting score ranges from -1 to 1, where a high score denotes that the data point fits well in the current cluster. We used the Louvain community detection algorithm implemented in “tl.louvain” of Scanpy package (**version 1.7.0**) for cell clustering. In our study, Louvain algorithm would generate much more cell clusters than real cell types when the resolution was 0.5 and far fewer when the resolution was 0.01. Thus, we set the resolution parameter range from 0.5 to 0.01 with a step of 0.01 and computed ARI score with *sklearn* package for each step. The maximum ARI score was employed as the final evaluation index. On account of BBKNN cannot give the corrected feature representation, we calculated the evaluation indexes in UMAP embedding space. The embedding was computed with the default parameters based on the same random seed through *umap-learn* package (**version 0.4.6**). For the *census of immune project*, we assessed the performance based on 20, 000 random sampled cells and averaged the scores of 10 replications.

### Cell marker inferring and cell type identification

The marker genes of cell clusters were calculated as mentioned in Miscell [13]. Specifically, we constructed a new deep neural network (denoted as *F: R*^*n*^ *-> [0,1]*) by freezing the parameters of the trained encoder and adding a single linear classifier at the end of it. The classifier was trained for cell cluster prediction. We used the importance score calculated by integrated gradient algorithm [43] as the surrogate metric for the impact of each gene on classification output. In specificity, the integrating gradient algorithm calculates the important score of the *i*^*th*^ gene as the gradient of *F(x)* along the *i*^*th*^ dimension, which is defined as:

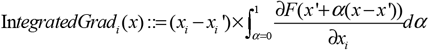

The *x* and *x’* are the actual and baseline expression levels respectively. We set *x’* to 0. A higher importance score represents a more significant impact of gene for the specific cell cluster. We manually annotated cell types according to genes with the highest importance scores.

### External software

Louvain community detection algorithm implemented in *Scanpy* package (**version 1.7.0**) was applied for cell clustering. We applied UMAP algorithm to visualize cells in a two-dimensional space if unspecified. UMAP failed on 3,424,607 cells after 72 hours; thus t-Distributed Stochastic Neighbor Embedding (t-SNE) algorithm from *FIt-SNE* package was utilized to construct a global view of HCA embedding space. Force-directed layout embedding (FLE) from Pegasus package was applied for trajectories inferring.

## Availability of data and materials

The source code of Fugue is available at https://github.com/xilinshen/Fugue. The datasets supporting the conclusions of this article are publicly available through onlines sources. The simulation dataset was available at https://github.com/xilinshen/Fugue/tree/master/data; the cell line dataset was available at https://support.10xgenomics.com/single-cell-gene-expression/datasets/1.1.0/jurkat, https://support.10xgenomics.com/single-cell-gene-expression/datasets/1.1.0/293t and https://support.10xgenomics.com/single-cell-gene-expression/datasets/1.1.0/jurkat:293t_50:50; the PBMC dataset was downloaded from https://www.ncbi.nlm.nih.gov/geo/query/acc.cgiãacc=GSE96583; all available single cells of HCA repository was downloaded from https://www.humancellatlas.org/.

## Author contribution

X.L., K.C. and L.S. designed and supervised the study. X.L. and X.S. wrote the manuscript. X.L., K.C., L.S. and X.S. revised the manuscript. H.S., D.W., M.F., J.H., J.L. collected the data. X.S., Y.Y., M.Y., Y.L. processed the data. X.S., X.L., K.C., L.S., Y.Y., M.Y. and Y.L. interpreted the results. All authors reviewed and approved the submission of this manuscript.

## Acknowledgment

We want to thank all the researchers for their generosity to make their data publicly available.

## Supplementary figure and table legends

**Supplementary Figure 1. Evaluation of Fugue’s robustness over changes of hyperparameters based on the *simulation dataset***. (A) Effect of momentum and queue size on the performance of Fugue. (B) The changes of loss values versus epochs. Error bands are standard deviations determined across 10 runs. (C) The performance of Fugue over the choice of data augmentation ratios.

**Supplementary Figure 2**. UMAP plot of batch effect removing performance on the *cell line dataset* across Seurat V3, ComBat, Harmony, BBKNN, Scanorama, scVI, Pegasus L/S adjustment and INSCT. Cells are colored by batch and cell type respectively.

**Supplementary Figure 3**. UMAP plot of batch effect removing performance on the *PBMC* dataset across Seurat V3, ComBat, Harmony, BBKNN, Scanorama, scVI, Pegasus L/S adjustment and INSCT. Cells are colored by batch and cell label.

**Supplementary Figure 4**. UMAP plot of the *census of immune project* before (A) and after (B-H) batch correction using ComBat, Harmony, BBKNN, Scanorama, scVI, Pegasus L/S adjustment and INSCT. Cells are colored by batch and cell type respectively.

**Supplementary Figure 5**. UMAP plot of batch effect removing performance on the *lung dataset* across Seurat V3, Harmony, ComBat, Scanorama, Scanorama, Pegasus L/S adjustment, scVI, BBKNN and INSCT. Cells are colored by batch.

**Supplementary Figure 6. Assessment of the performance of Fugue on the *brain dataset***. (A-b) UMAP plot showing cells in the *brain dataset* before (A) and after (B-C) Fugue integration. Cells are colored by batch in (A-B) and cell cluster label in (C). (D) Bar plot depicting kBET scores of different batch effect removing methods on HCA brain cohorts. (E) Expression of cell type markers across the integrated embedding space. Dark and light colors represent low and high relative expression values, respectively. (F) Dot plot of cell type markers across batches. The size of each circle reflects the percentage of cells in a cluster where the gene is detected, and the color intensity reflects the average expression level within each cluster.

**Supplementary Figure 7. The similarity across batch embedding representation of all samples in HCA repository**. (A) Heatmap of cosine similarity of dimension reduction representations of the batch embedding matrix across all samples. Each red frame represents samples from one cohort. (B-D) show 3 representative projects from (A), namely C2, C1 and C39.

**Supplementary Figure 8. Fugue inferred cell clusters from HCA embedding space**. (A) Importance scores of the top 5 marker genes for each cell cluster. Representative markers are displayed on the right side. (B) Bar plot displaying the cohort composition of cell clusters.

**Supplementary Figure 9**. TSNE plot of all quality-controlled cells from HCA. TSNE plot in the top left corner is labeled by cell cluster labels. The others are colored by the expression level of marker genes of immune cells. Light and deep red represent low and high relative expression values, respectively.

**Supplementary Figure 10**. Violin plot deciphering expression levels of cell type markers across immune cell subtype in HCA repository. Violins were colored by cohorts. Cohorts with more than 1000 cells were displayed.

**Supplementary Figure 11**. Dot plot of cell type markers of cardiac cells across batches. The size of each circle reflects the percentage of cells in a cluster where the gene is detected, and the color intensity reflects the average expression level within each cluster.

**Supplementary Figure 12**. Correlation of pseudo-time and expression level of cell differentiation markers across cardiac cell types. The curves representing polynomial fits for each batch.

**Supplementary Figure 13. FLE embedding space of the *census of immune project* integrated by Fugue**. Cells are colored by batch (A) and cell type (B-C). (B) and (C) displaying the major cell types in cord blood (B) and bone marrow (C), respectively.

**Supplementary Figure 14. UMAP plot of the simulated cells**. (A-B) deciphering the *simulation dataset* and (C-D) deciphering the *simulation_rm dataset*, which was obtained by manually removing cell type 1 from batches 2-5 and retaining them in batch 1.

**Supplementary Table 1**. Detailed information of datasets from HCA repository.

**Supplementary Table 2**. The 250 genes with the highest importance scores of HCA cell clusters were inferred from Fugue. Marker genes of each cluster were colored in blue.

**Supplementary Table 3**. Detailed information of benchmark datasets, their gene filtering and hyperparameter settings of Fugue.

**Supplementary Table 4**. Detailed information of the benchmark methods.

