## Supplementary Figure 1-14 and Supplementary Table 4 for "Scalable batch-correction approach for integrating large-scale single-cell transcriptomes"

### Supplementary Figures

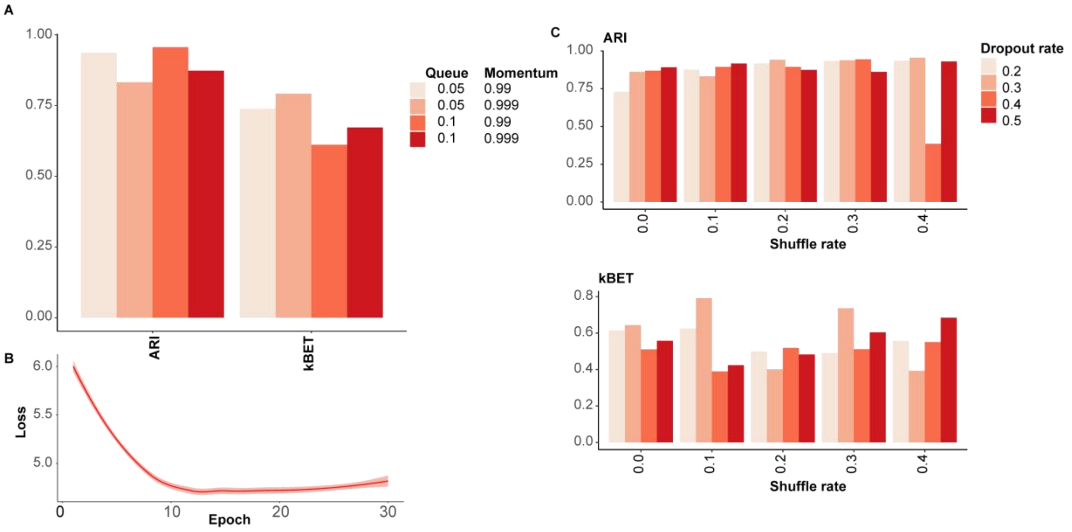


**Supplementary Figure 1. Evaluation of Fugue's robustness over changes of hyperparameters based on the *simulation dataset*.** (A) Effect of momentum and queue size on the performance of Fugue. (B) The changes of loss values versus epochs. Error bands are standard deviations determined across 10 runs. (C) The performance of Fugue over the choice of data augmentation ratios.

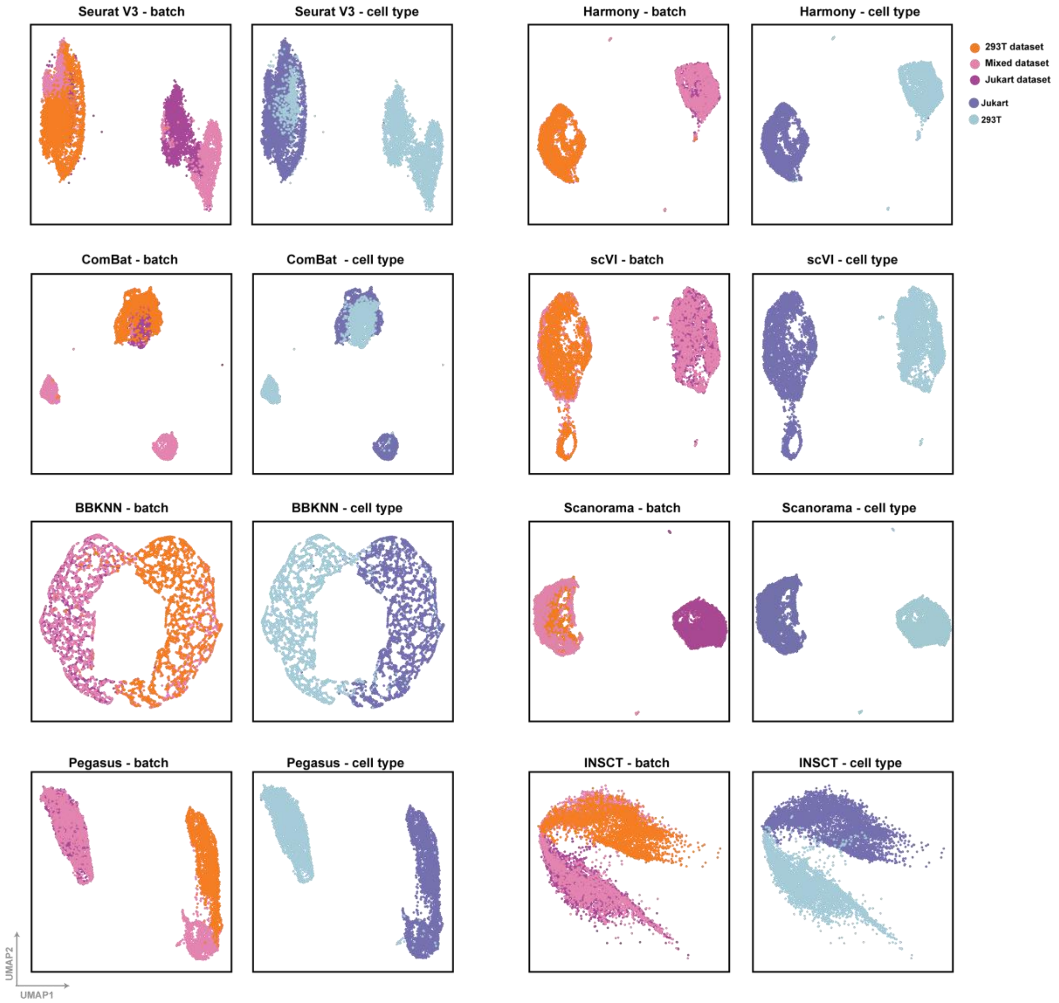


**Supplementary Figure 2.** UMAP plot of batch effect removing performance on the *cell line dataset* across Seurat V3, ComBat, Harmony, BBKNN, Scanorama, scVI, Pegasus L/S adjustment and INSCT. Cells are colored by batch and cell type respectively.

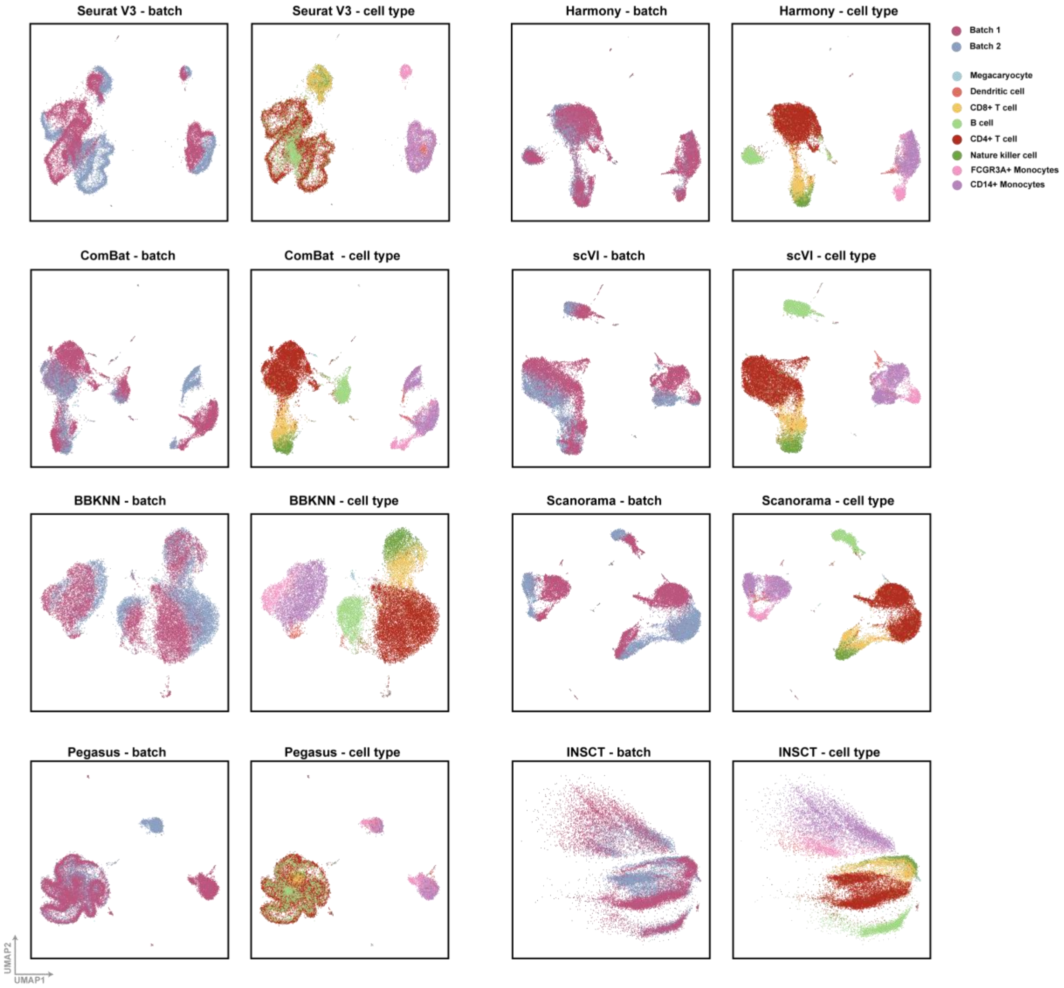


**Supplementary Figure 3.** UMAP plot of batch effect removing performance on the *PBMC* dataset across Seurat V3, ComBat, Harmony, BBKNN, Scanorama, scVI, Pegasus L/S adjustment and INSCT. Cells are colored by batch and cell label.

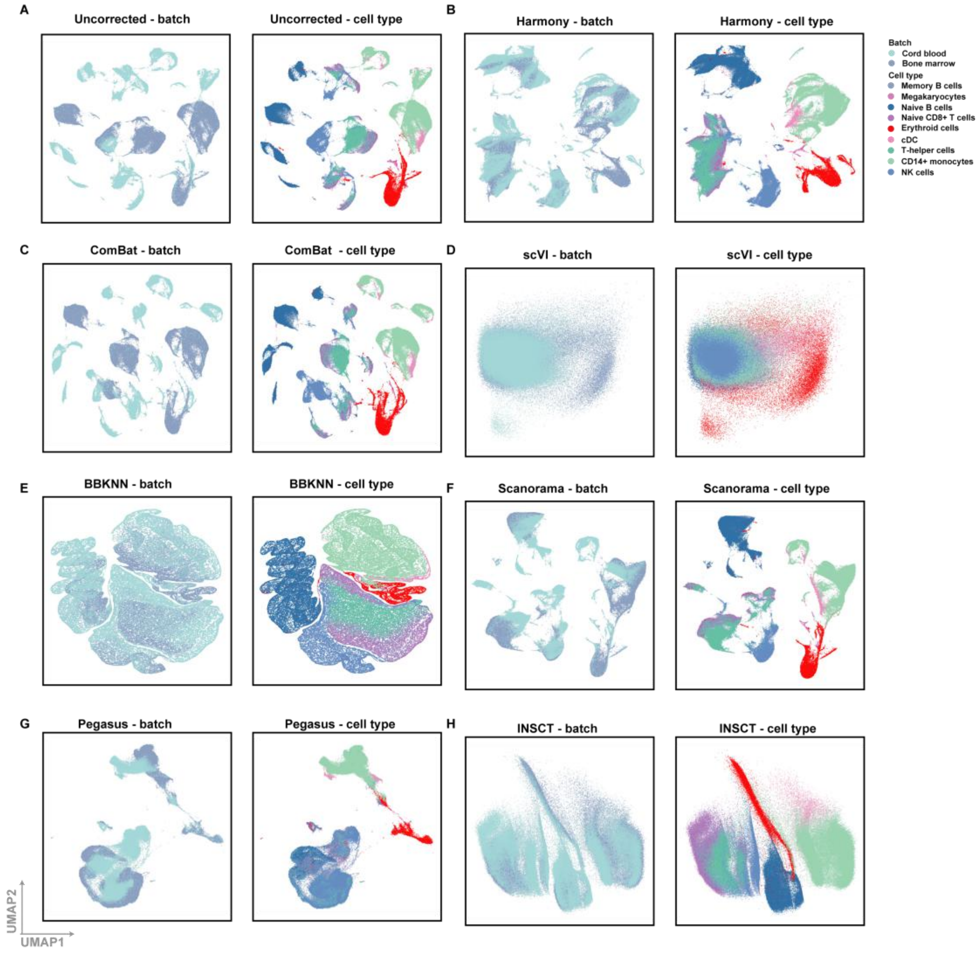


**Supplementary Figure 4.** UMAP plot of the *census of immune project* before (A) and after (B-H) batch correction using ComBat, Harmony, BBKNN, Scanorama, scVI, Pegasus L/S adjustment and INSCT. Cells are colored by batch and cell type respectively. cDC, classical dendritic cell.

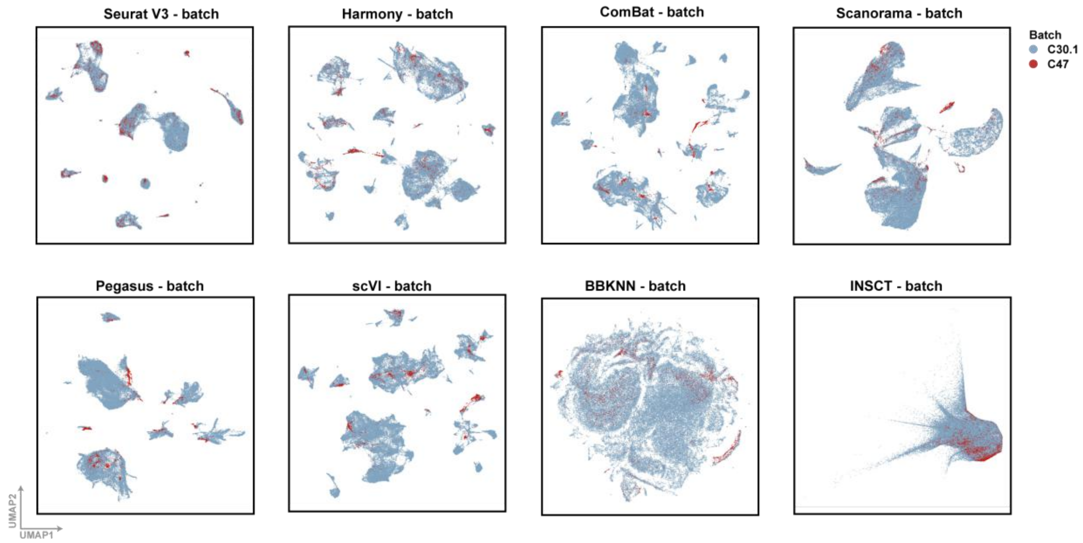


**Supplementary Figure 5.** UMAP plot of batch effect removing performance on the *lung dataset* across Seurat V3, Harmony, ComBat, Scanorama, Scanorama, Pegasus L/S adjustment, scVI, BBKNN and INSCT. Cells are colored by batch.

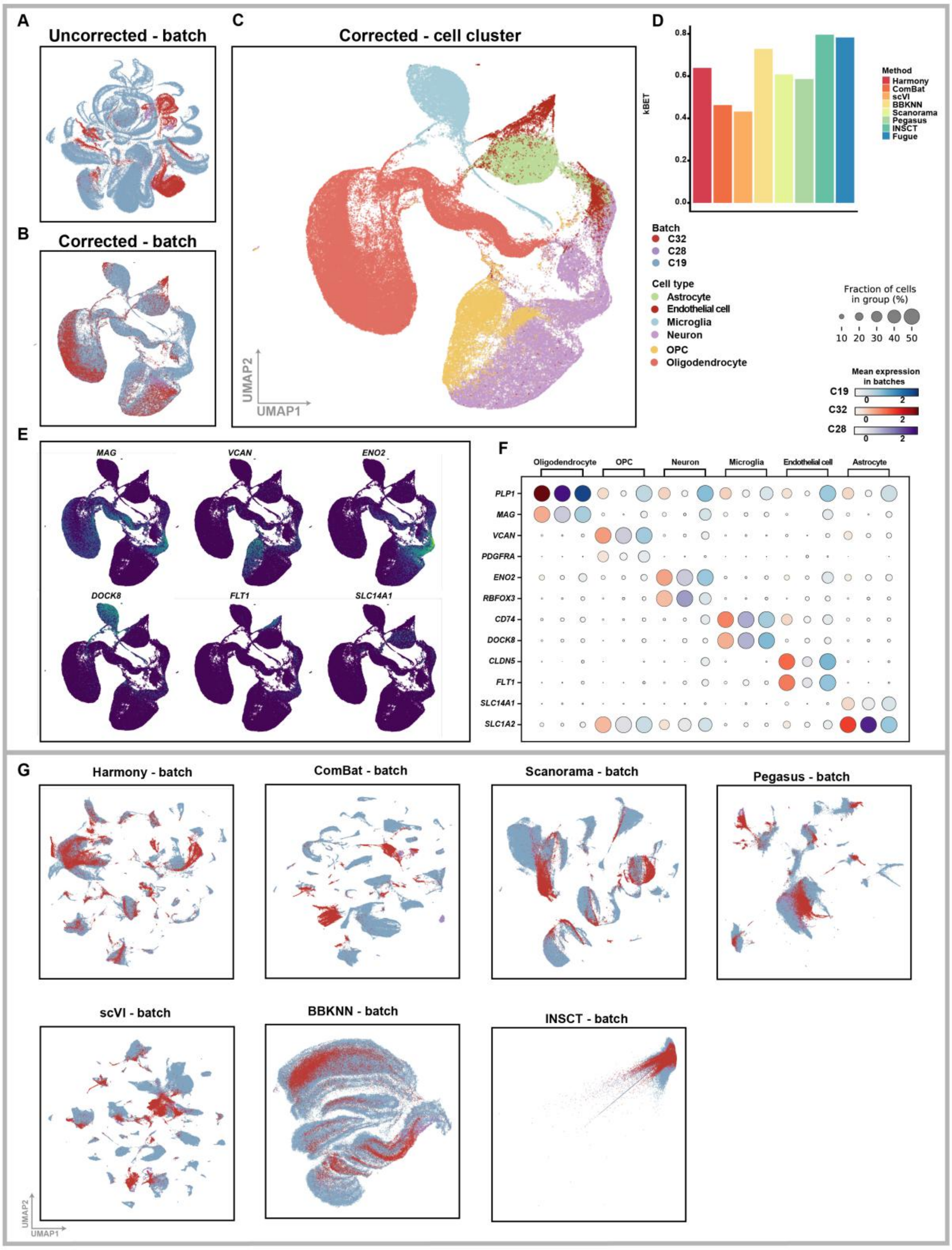


**Supplementary Figure 6. Assessment of the performance of Fugue on the *brain dataset*.** (A-b) UMAP plot showing cells in the *brain dataset* before (A) and after (B-C) Fugue integration. Cells are colored by batch in (A-B) and cell cluster label in (C). (D) Bar plot depicting kBET scores of different batch effect removing methods on HCA brain cohorts. (E) Expression of cell type markers across the integrated embedding space. Dark and light colors represent low and high relative expression values, respectively. (F) Dot plot of cell type markers across batches. The size of each circle reflects the percentage of cells in a cluster where the gene is detected, and the color intensity reflects the average expression level within each cluster. OPC, oligodendrocyte progenitor cell.

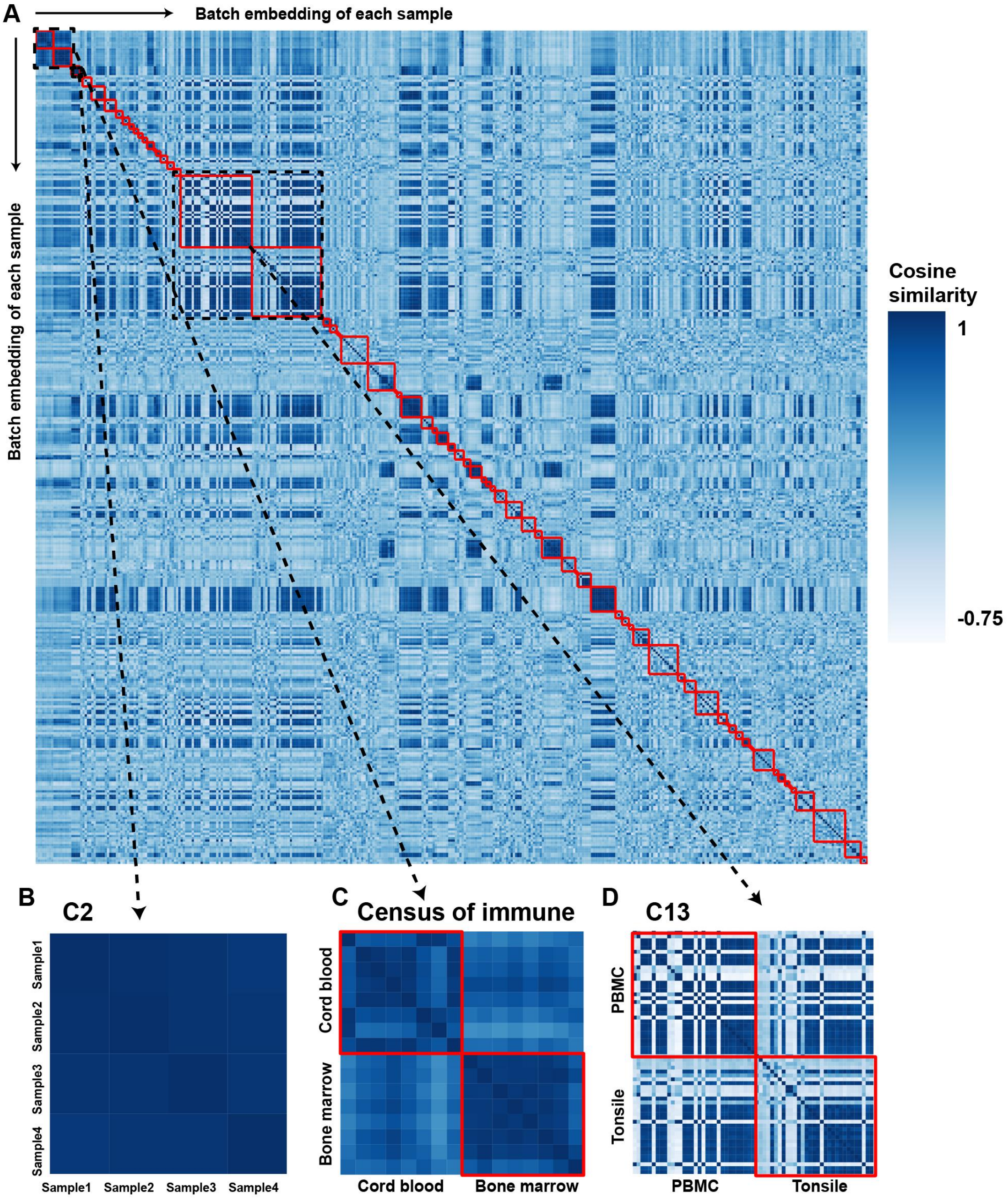


**Supplementary Figure 7. The similarity across batch embedding representation of all samples in HCA repository.** (A) Heatmap of cosine similarity of dimension reduction representations of the batch embedding matrix across all samples. Each red frame represents samples from one cohort. (B-D) show 3 representative projects from (A), namely C2, C1 and C39.

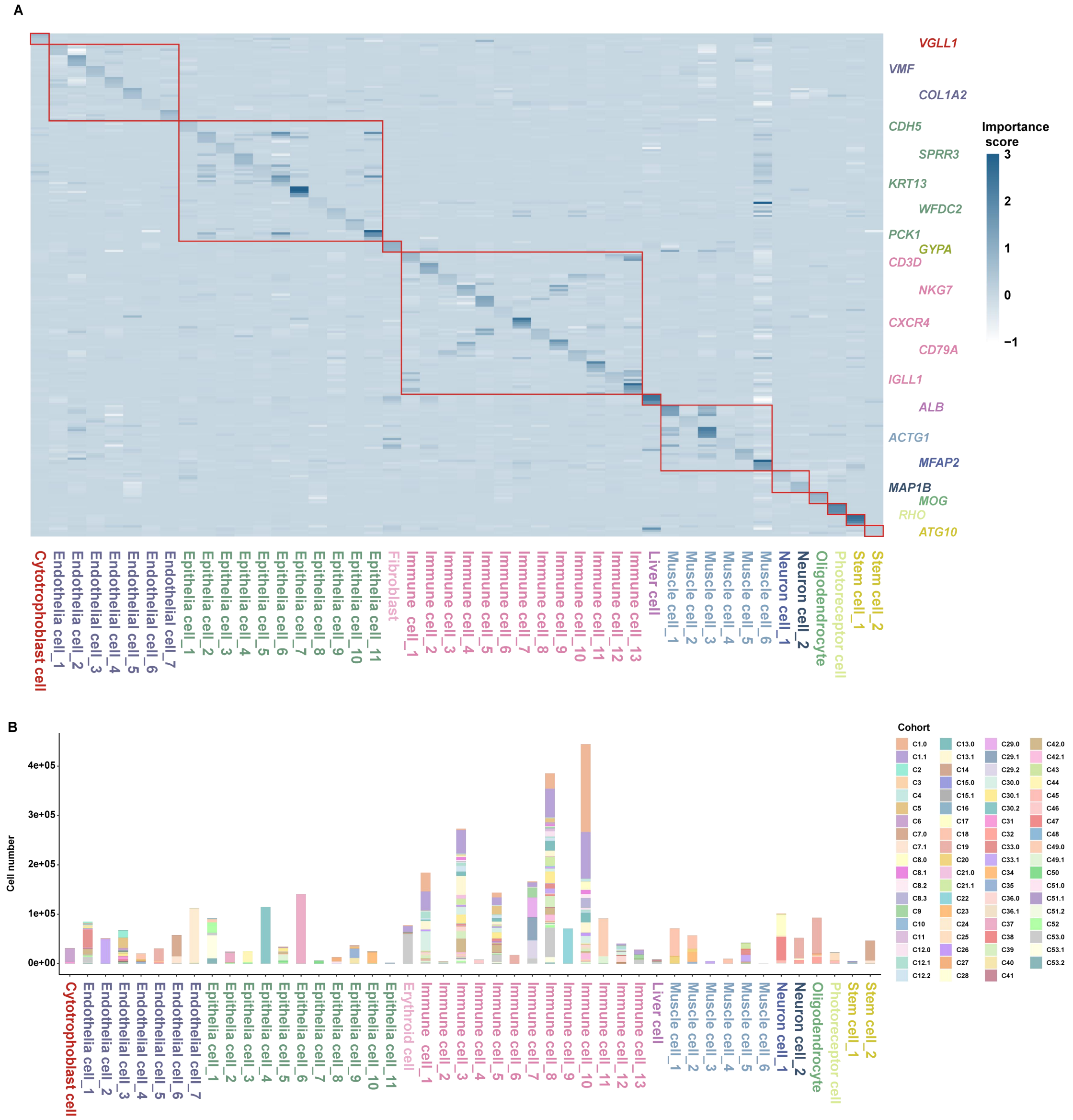


**Supplementary Figure 8. Fugue inferred cell clusters from HCA embedding space.** (A) Importance scores of the top 5 marker genes for each cell cluster. Representative markers are displayed on the right side. (B) Bar plot displaying the cohort composition of cell clusters.

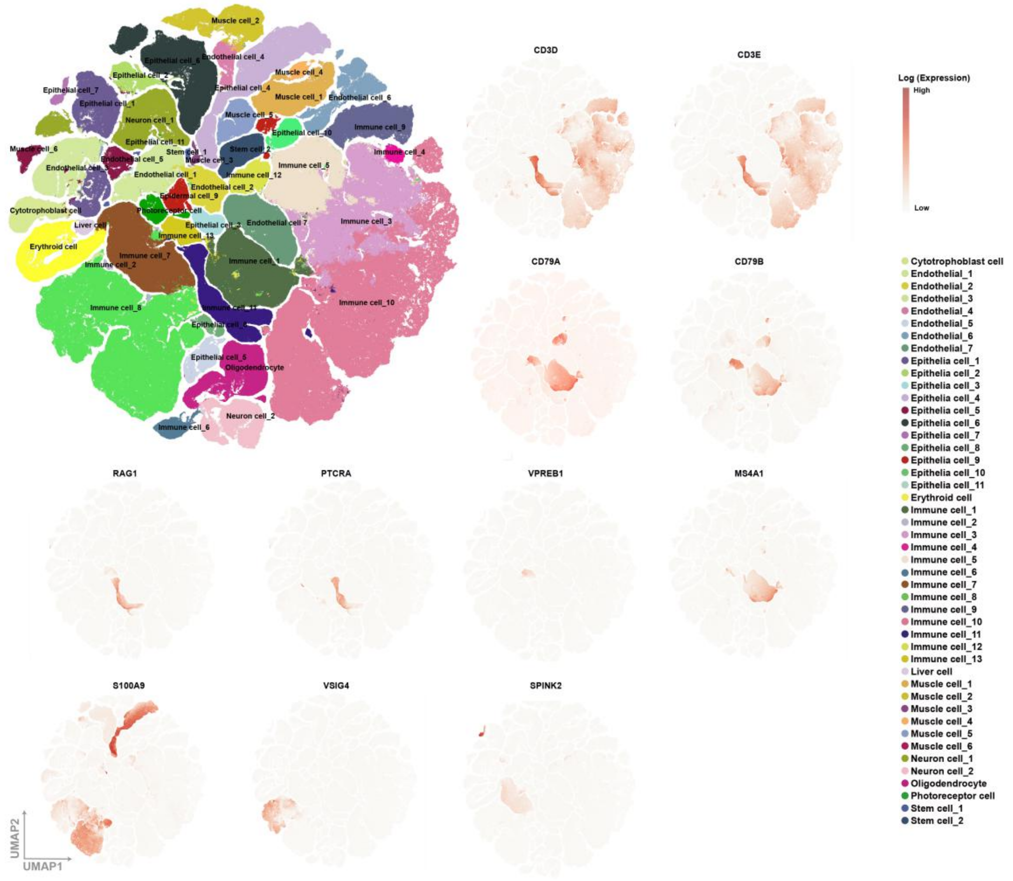


**Supplementary Figure 9.** TSNE plot of all quality-controlled cells from HCA. TSNE plot in the top left corner is labeled by cell cluster labels. The others are colored by the expression level of marker genes of immune cells. Light and deep red represent low and high relative expression values, respectively.

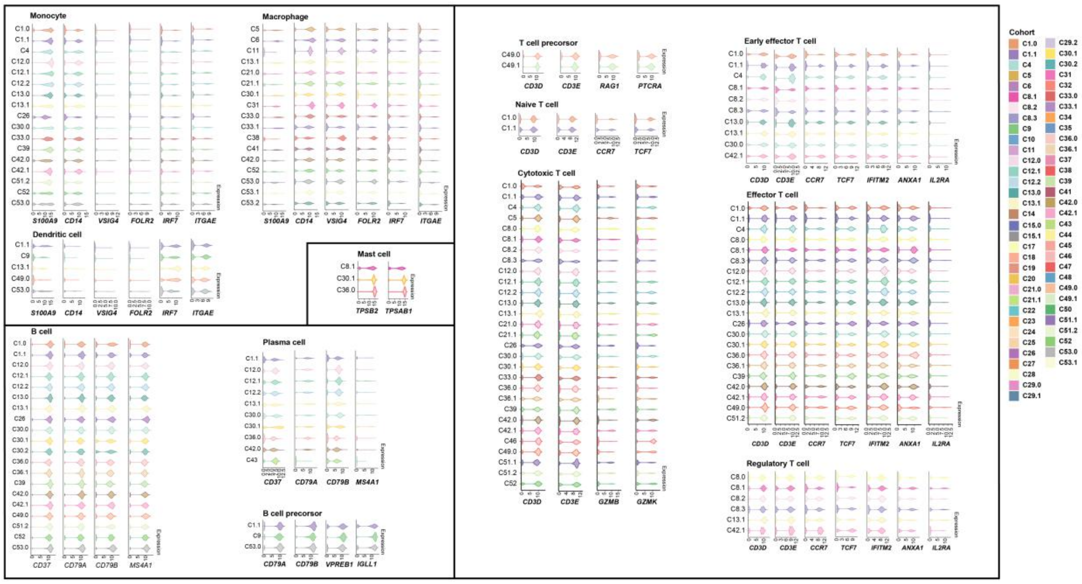


**Supplementary Figure 10.** Violin plot deciphering expression levels of cell type markers across immune cell subtype in HCA repository. Violins were colored by cohorts. Cohorts with more than 1000 cells were displayed.

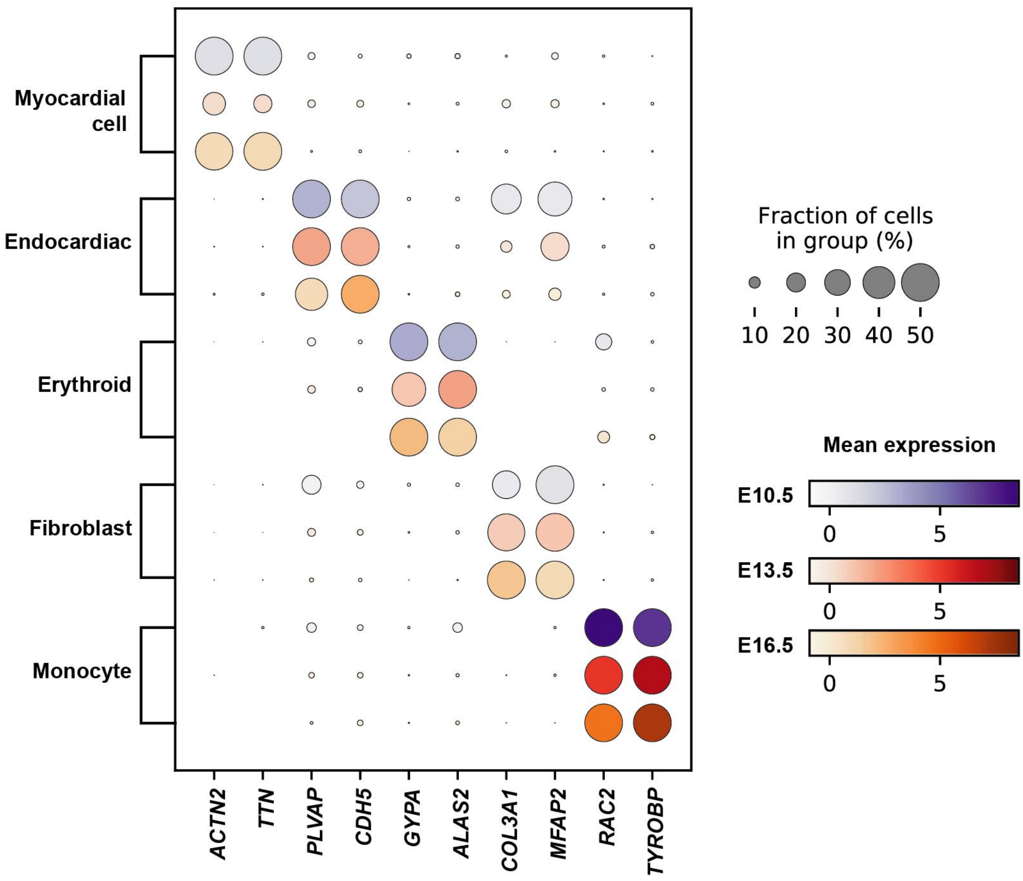


**Supplementary Figure 11.** Dot plot of cell type markers of cardiac cells across batches. The size of each circle reflects the percentage of cells in a cluster where the gene is detected, and the color intensity reflects the average expression level within each cluster.

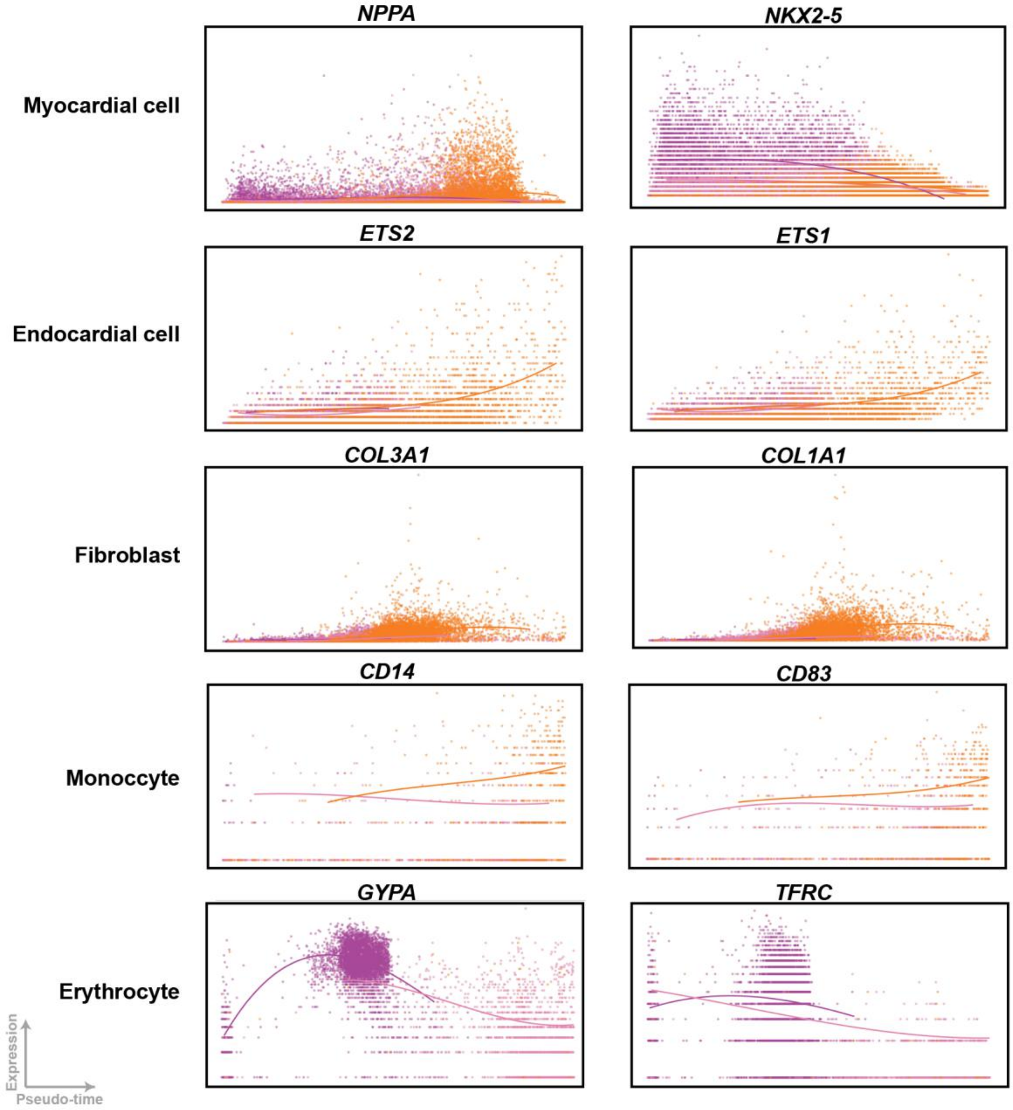


**Supplementary Figure 12.** Correlation of pseudo-time and expression level of cell differentiation markers across cardiac cell types.

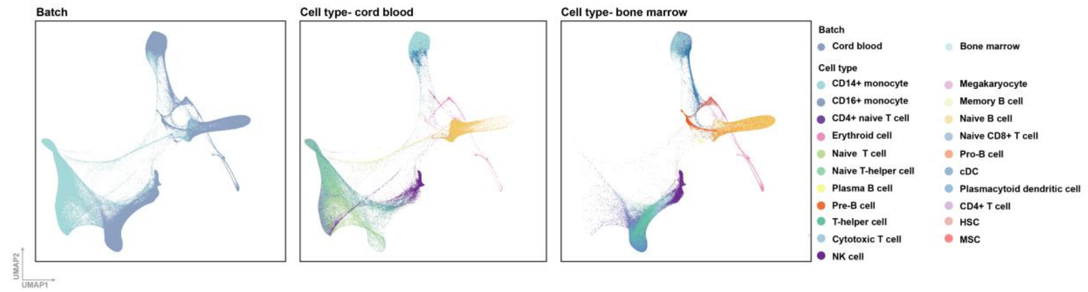


**Supplementary Figure 13. FLE embedding space of the *census of immune project* integrated by Fugue.** Cells are colored by batch (A) and cell type (B-C). (B) and (C) displaying the major cell types in cord blood (B) and bone marrow (C), respectively. NK cell, natural killer cell; cDC, classical dendritic cell; HSC, hematopoietic stem progenitor cell; MSC, multipotent progenitor cell.

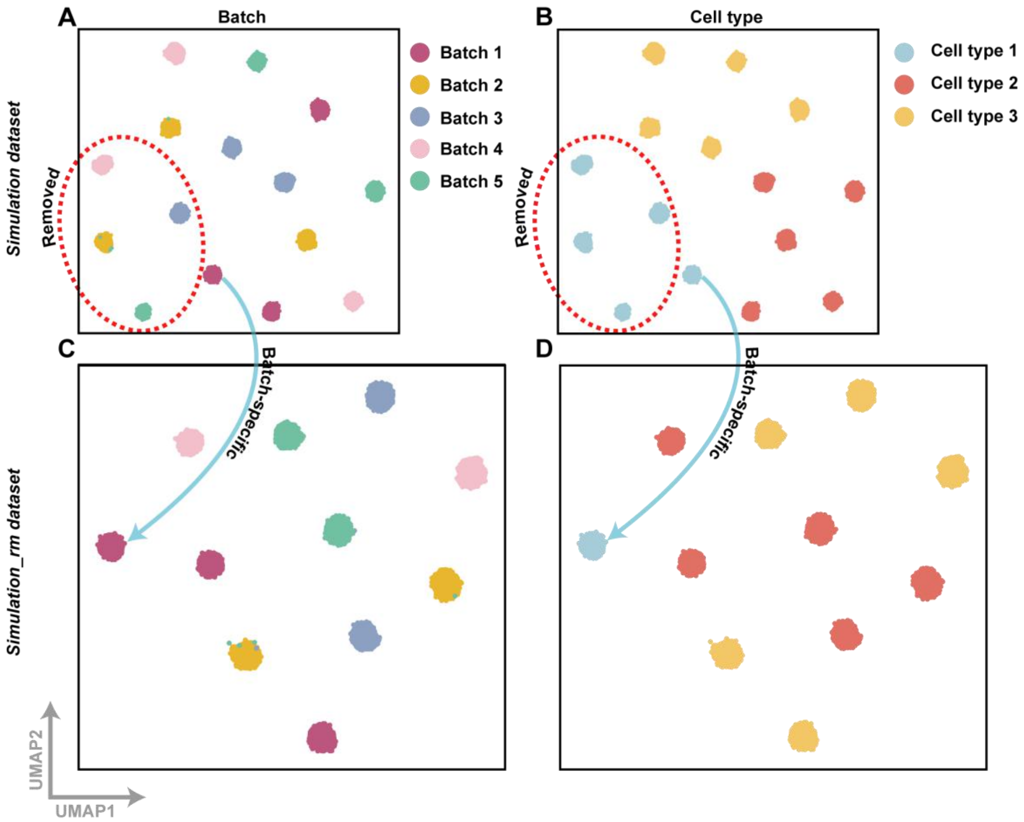


**Supplementary Figure 14. UMAP plot of the simulated cells.** (A-B) deciphering the *simulation dataset* and (C-D) deciphering the *simulation_rm dataset,* which was obtained by manually removing cell type 1 from batches 2-5 and retaining them in batch 1.

### Supplementary Tables

**Supplementary Table 1.** Detailed information of datasets from HCA repository.

**Supplementary Table 2.** The 250 genes with the highest importance scores of HCA cell clusters were inferred from Fugue. Marker genes of each cluster were colored in blue.

**Supplementary Table 3.** Detailed information of benchmark datasets, their gene filtering and hyperparameter settings of Fugue.

**Supplementary Table 4.** Detailed information of the benchmark methods.

| Method | Package | Environment | Version | DOI |
|---|---|---|---|---|
| Seurat V3 | seurat | R 3.6 | 3.2.0 | 10.1038/nbt.4096 |
| Harmony | pegasus | Python 3.8 | 1.40 | s41592-019-0619-0 |
| ComBat | scanpy | Python 3.8 | 1.7.0 | 10.1093/bioinformatics/bts034 |
| scVI | scvi | Python 3.8 | 0.0.0 | s41592-018-0229-2 |
| BBKNN | scanpy | Python 3.8 | 1.7.0 | 10.1093/bioinformatics/btz625 |
| Scanorama | scanorama | Python 3.8 | 1.70 | s41587-019-0113-3 |
| Pegasus | pegasus | Python 3.8 | 1.40 | s41592-020-0905-x |
| INSCT | tnn | Python 3.8 | 0.0.2 | s42256-021-00361-8 |
